# Dhaka: Variational Autoencoder for Unmasking Tumor Heterogeneity from Single Cell Genomic Data

**DOI:** 10.1101/183863

**Authors:** Sabrina Rashid, Sohrab Shah, Ziv Bar-Joseph, Ravi Pandya

**Affiliations:** Computational Biology Department, Carnegie Mellon University, Pittsburgh, USA; Department of Computer Science, University of British Columbia, Vancouver, Canada; Department of Molecular Oncology, BC Cancer Agency, Vancouver, Canada; Department of Pathology and Laboratory Medicine, University of British Columbia, Vancouver, Canada; Machine Learning Department and Computational Biology Department, Carnegie Mellon University, Pittsburgh, USA; Microsoft Research, Redmond, USA

**Keywords:** Single cell genomic data, Neural Networks, Tumor Heterogeneity, Differentiation and/or lineage trajectory

## Abstract

**Motivation:** Intra-tumor heterogeneity is one of the key confounding factors in deciphering tumor evolution. Malignant cells exhibit variations in their gene expression, copy numbers, and mutation even when originating from a single progenitor cell. Single cell sequencing of tumor cells has recently emerged as a viable option for unmasking the underlying tumor heterogeneity. However, extracting features from single cell genomic data in order to infer their evolutionary trajectory remains computationally challenging due to the extremely noisy and sparse nature of the data.

**Results:** Here we describe ‘Dhaka’, a variational autoencoder method which transforms single cell genomic data to a reduced dimension feature space that is more efficient in differentiating between (hidden) tumor subpopulations. Our method is general and can be applied to several different types of genomic data including copy number variation from scDNA-Seq and gene expression from scRNA-Seq experiments. We tested the method on synthetic and 6 single cell cancer datasets where the number of cells ranges from 250 to 6000 for each sample. Analysis of the resulting feature space revealed subpopulations of cells and their marker genes. The features are also able to infer the lineage and/or differentiation trajectory between cells greatly improving upon prior methods suggested for feature extraction and dimensionality reduction of such data.

**Availability and Implementation:** All the datasets used in the paper are publicly available and developed software package is available on Github https://github.com/MicrosoftGenomics/Dhaka.

Supporting info and Software: https://github.com/MicrosoftGenomics/Dhaka

## Introduction

Tumor cells are often very heterogeneous. Typical cancer progression consists of a prolonged clinically latent period during which several new mutations arise leading to changes in gene expression and DNA copy number for several genes de Bruin et al. (2014), Andor et al. (2016), Min et al. (2015). As a result of such genomic variability, we often see multiple subpopulations of cells within a single tumor.

The goal of effective cancer treatment is to treat all malignant cells without harming the originating host tissue. Clinical approaches should thus take into account the underlying evolutionary structure in order to identify treatments that can specifically target malignant cells while not affecting their normal cell of origin. It is also important to determine if the ancestral tumor clones eventually disappear (chain like evolution) or if several genotypically different clones of cells evolved in parallel (branched evolution) de Bruin et al. (2014). Tumors resulting from these two evolutionary trajectories respond differently and ignoring the evolutionary process when determining treatment can lead to therapy resistance and possible cancer recurrence. Thus, characterization of the hidden subpopulations and their underlying evolutionary structure is an important issue for both the biological understanding and clinical treatment of cancer. Prior studies have mainly relied on bulk sequencing to investigate tumor evolution Navin and Hicks (2010), Russnes et al. (2011). In such experiments thousands of cells are sequenced together, which averages out the genomic characteristics of the individual cells making it hard to infer these sub-populations. More recently, single cell sequencing has emerged as a useful tool to study such cellular heterogeneity Venteicher et al. (2017), Tirosh et al. (2016b), Zahn et al. (2017), Giustacchini et al. (2017).

While single cell data is clearly much more appropriate for addressing tumor heterogeneity and evolution, it also raises new computational and experimental challenges. Due to technical challenges (for example, the low quantity of genetic material and the coverage for each of the cells sequenced) the resulting data is often very noisy and sparse with many dropout events Gawad et al. (2016), Zong et al. (2012). These issues affect both, scRNA-Seq and scDNA-Seq experiments which are used for copy number and mutation estimation. Given these issues, it remains challenging to identify meaningful features that can accurately characterize the single cells in terms of their clonal identity and differentiation state. To address this, several methods have been proposed to transform the observed gene expression or copy number profiles in order to generate features that are more robust for downstream analysis. However, as we show below, many of the feature transformation techniques that are usually applied to genomic data fail to identify the sub populations and their trajectories. For example, while t-SNE Maaten and Hinton (2008) and diffusion maps Roweis and Saul (2000) are very successful in segregating cells between different tumor samples, they are less successful when trying to characterize the evolutional trajectories of a single tumor. Recently, several unsupervised feature transformation were proposed for analysis of single-cell RNA-seq data Pierson and Yau (2015), Wang et al. (2017), Li et al. (2017), DeTomaso and Yosef (2016). Among these tools, ZIFA Pierson and Yau (2015) explicitly models the dropout event in single cell RNA seq data to improve the reduced dimension representation whereas SIMLR Wang et al. (2017) developed a new similarity learning framework that can be used in conjunction with t-SNE to reduce dimension of the data. In addition to dimensionality reduction methods several single cell clustering algorithm has been proposed as well Xu and Su (2015), Fan et al. (2016). SNN-cliq Xu and Su (2015) constructs a shared k-nearest neighbor graph across all cells and then finds maximal cliques and PAGODA relies on prior set of annotated genes to find transcriptomal heterogeneity. All these methods can successfully distinguish between different groups of cells in a dataset. However, such methods are not designed for determining the relationship between the detected clusters which is the focus of tumor evolutionary analysis. In addition, most current single cell clustering methods are focused on only one type of genomic data (for example scRNA-Seq) and do not work well for multiple types of such data.

Another direction that has been investigated for reducing the dimensionality of scRNA-Seq data is the use of neural networks (NN) Lin et al. (2017), Gupta et al. (2015). In Lin *et al*. Lin et al. (2017), the authors used prior biological knowledge including protein-protein and protein-DNA interaction to learn the architecture of a NN and subsequently project the data to a lower dimensional feature space. Unlike these prior approaches, which were supervised, we are using neural networks in a completely unsupervised manner and so do not require labeled data as prior methods have. Specifically, in our software ‘Dhaka’ we have developed a variational autoencoder that combines Bayesian inference with unsupervised deep learning, to learn a probabilistic encoding of the input data. Our autoencoder method can be used for analyzing different types of genomic data. Specifically, in this paper we have analyzed 4 scRNA-Seq and 2 scDNA-Seq datasets. We used the variational autoencoder to project the expression and copy number profiles of tumor populations and were able to capture clonal evolution of tumor samples even for noisy sparse datasets with very low coverage. We also compare the performance of Dhaka with two generalized dimensionality reduction methods, PCA Jolliffe (1986) and t-SNE Maaten and Hinton (2008) and two specialized single cell dimensionality methods ZIFA Pierson and Yau (2015) and SIMLR Wang et al. (2017). Dhaka shows significant improvement over the prior methods thus corroborates the effectiveness of our method in extracting important biological and clinical information from cancer samples.

## Methods

### Variational autoencoder

We used a variational autoencoder to analyze single cell genomic data. For this, we adapted an autoencoder initially proposed by Kingma and Welling (2013). *Autoencoders* are multilayered perceptron neural networks that sequentially deconstruct data (x) into latent representation (z) and then use these representations to reconstruct outputs that are similar (in some metric space) to the inputs. The main advantage of this approach is that the model learns the best features and input combinations in a completely unsupervised manner. In *variational autoencoders (VAE)* unsupervised deep learning is combined with Bayesian inference. Instead of learning an unconstrained representation of the data we impose a regularization constraint. We assume that the latent representation is coming from a probability distribution, in this case a multivariate Gaussian (*N*(*μ_z_, σ_z_*)). The intuition behind such representation for single cell data is that the heterogeneous cells are actually the result of some underlying biological process leading to the observed expression and copy number data. These processes are modeled here as distribution over latent space, each having their distinct means and variances. Hence the autoencoder actually encodes not only the means (*μ_z_*) but also the variances (*σ_z_*) of the Gaussian distributions. The latent representation (*z*) is then sampled from the learned posterior distribution *qϕ*(*z*|*x*) ~ N(*μ_z_, σ_z_I*). Here *ϕ* are the parameters of the encoder network (such as biases and weights). The sampled latent representation is then passed through a similar decoder network to reconstruct the input *x̄* ~ *ρ_θ_*(*x*|*z*), where *θ* are the parameters of the decoder network. Although the model is trained holistically, we are actually interested in the latent representation *z* of the data since it represents the key information needed to accurately reconstruct the inputs.

#### Model structure

Fig. 1 presents the structure of the autoencoder used in this paper. The input layer consists of nodes equal to the number of genes we are analyzing for each cell. We have used Rectified Linear unit(ReLu) activation function in all the layers except the final layer of getting the reconstructed output. We used sigmoid activation function in the final layer. We have used three intermediate layers with 1024, 512, and 256 nodes and a 3-dimensional latent layer. The latent layer has three nodes for mean (*μz*) and three nodes for variance (*σ_z_*), which in turn generate the 3 dimensional latent variable *z*.

**Fig. 1.**
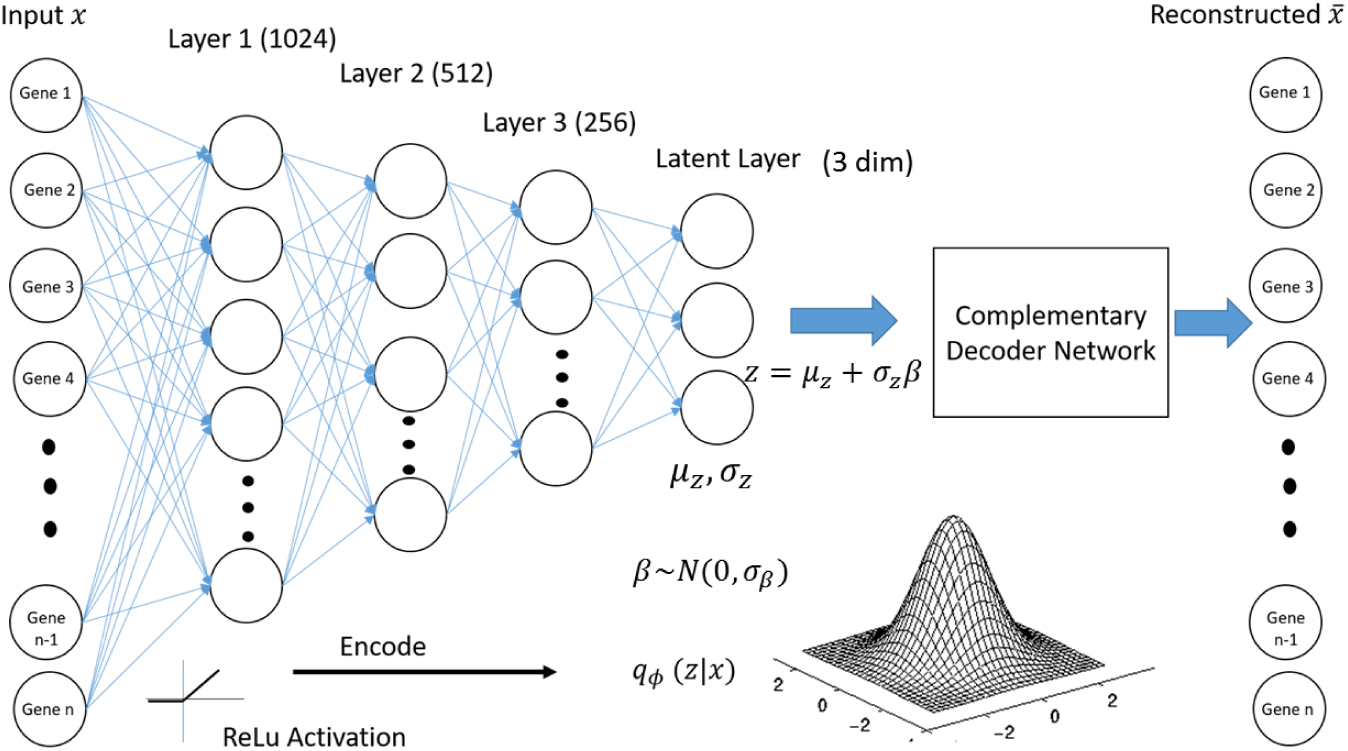
Structure of the variational autoencoder used in Dhaka.

The size of the latent dimension (i.e. the representation we extract from the model) is a parameter of the model. As we show in Results, for the data analyzed in this paper three latent variables are enough to obtain accurate separation of cell states for both the expression and copy number datasets. Increasing this number did not improve the results and so all figures and subsequent analysis are based on this number. However, the method is general and if needed can use more or less nodes in the latent layer.

All datasets we analyzed had more than 5K genes and the reported structure with atleast 1024 nodes in the first intermediate layer (Fig. 1) was sufficient for them. We used three intermediate layers to gradually compress the encoding to a 3 dimensional feature space. We have also compared three different structures of autoencoders: i) the proposed three intermediate layers, ii) one intermediate layer, and iii) five intermediate layers in the Results section.

#### Learning

To learn the parameters of the autoencoder, *ϕ* and *θ*, we need to maximize *log*(*p*(*x*|*ϕ, θ*)), the log likelihood of the data points *x*, given the model parameters. The marginal likelihood *log*(*p*(*x*)) is the sum of a variational lower bound Kingma and Welling (2013) and the Kullback-Leibler (KL) Joyce (2011) divergence between the approximate and true posteriors.

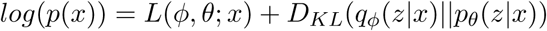

The likelihood L can be decomposed as following:

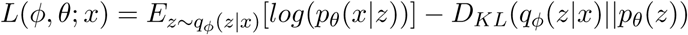

The first term can be viewed as the typical reconstruction loss intrinsic to all autoencoders, the second term can be viewed as the penalty for forcing the encoded representation to follow the Gaussian prior (the regularization part). We then use ‘RMSprop’, which relies on a variant of stochastic minibatch gradient descent, to minimize—*L*. In ‘RMSprop’, the learning rate weight is divided by the running average of the magnitudes of recent gradients for that weight leading to better convergence Tieleman and Hinton (2012). Detailed derivation of the loss computation can be found in Kingma and Welling (2013). To demonstrate the robustness of the training, we have shown the loss function plot from 50 independent trials on the Oligodendroglioma dataset (Supporting Fig. S1). The low standard error in the plot corroborates the robustness of training in Dhaka.

An issue in learning VAE with standard gradient descent is that gradient descent requires the model to be differentiable, which is inherently deterministic. However, in VAE the fact that we sample from the latent layer makes the model stochastic. To enable the use of gradient descent in our model, we introduce a new random variable *β*. Instead of sampling *z* directly from the *N* (*μ_z_, σ_z_I*), we set

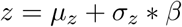

Where *β* is the Gaussian noise, *β* ~ *N*(0, *σ_β_*). Using *β* we do not need to sample from the latent layer and so the model is differentiable and gradient descent can be used to learn model parameters LeCun et al. (2015). *σ_β_* is the standard deviation of the Gaussian noise and is an input parameter of the model.

## Results

### Simulated dataset

We first performed simulation analysis to compare the Dhaka method with prior dimensionality reduction methods that have been extensively used for scRNA-Seq data: t-SNE Maaten and Hinton (2008), PCA Jolliffe (1986), ZIFA Pierson and Yau (2015), and SIMLR Wang et al. (2017). We generated a simulated dataset with 3K genes and 500 cells. To generate the simulated data we followed the method described in Yu et al. (2017). In the simulated dataset, cells are generated from five different clusters with 100 cells each. We simulate a total of 3000 genes in this dataset. All the 3000 genes are sampled with variable amounts of noise. Of these, 500 genes are sampled based on cluster specific expression profile while the expression of the remaining 2500 is sampled from a null model, i.e., completely noisy. We have used a Gaussian Mixture Model to cluster the reduced dimension data obtained from Dhaka and other competing methods and Bayesian Information criterion (BIC) to select the number of clusters. We next compute the Adjusted Rand Index (ARI) metric to determine the quality of resulting clustering for each dimensionality reduction method. Fig 2 shows the result of our autoencoder, PCA, t-SNE, ZIFA, and SIMLR projection for the 2500 noisy genes simulated data. As can be seen, the Dhaka autoencoder has the highest ARI score of 0.73. The closest is SIMLR (ARI: 0.70) and the ZIFA (ARI: 0.58). Although the Dhaka autoencoder identifies 4 clusters compared to SIMLR identifying 5, the cluster labels are better preserved in Dhaka autoencoder leading to higher ARI score. Implementation details of the competing methods can be found in Appendix 1.6. The variational autoencoder in Dhaka is optimized with no guaranteed global convergence. Hence we will see slightly different outputs with each run of the algorithm. We analyzed the robustness of the method to random initializations on the simulated dataset. With 10 random initializations we observed mean ARI of 0.73 with standard error of 0.01. This relatively low standard error corroborates the robustness of the proposed method.

**Fig. 2.**
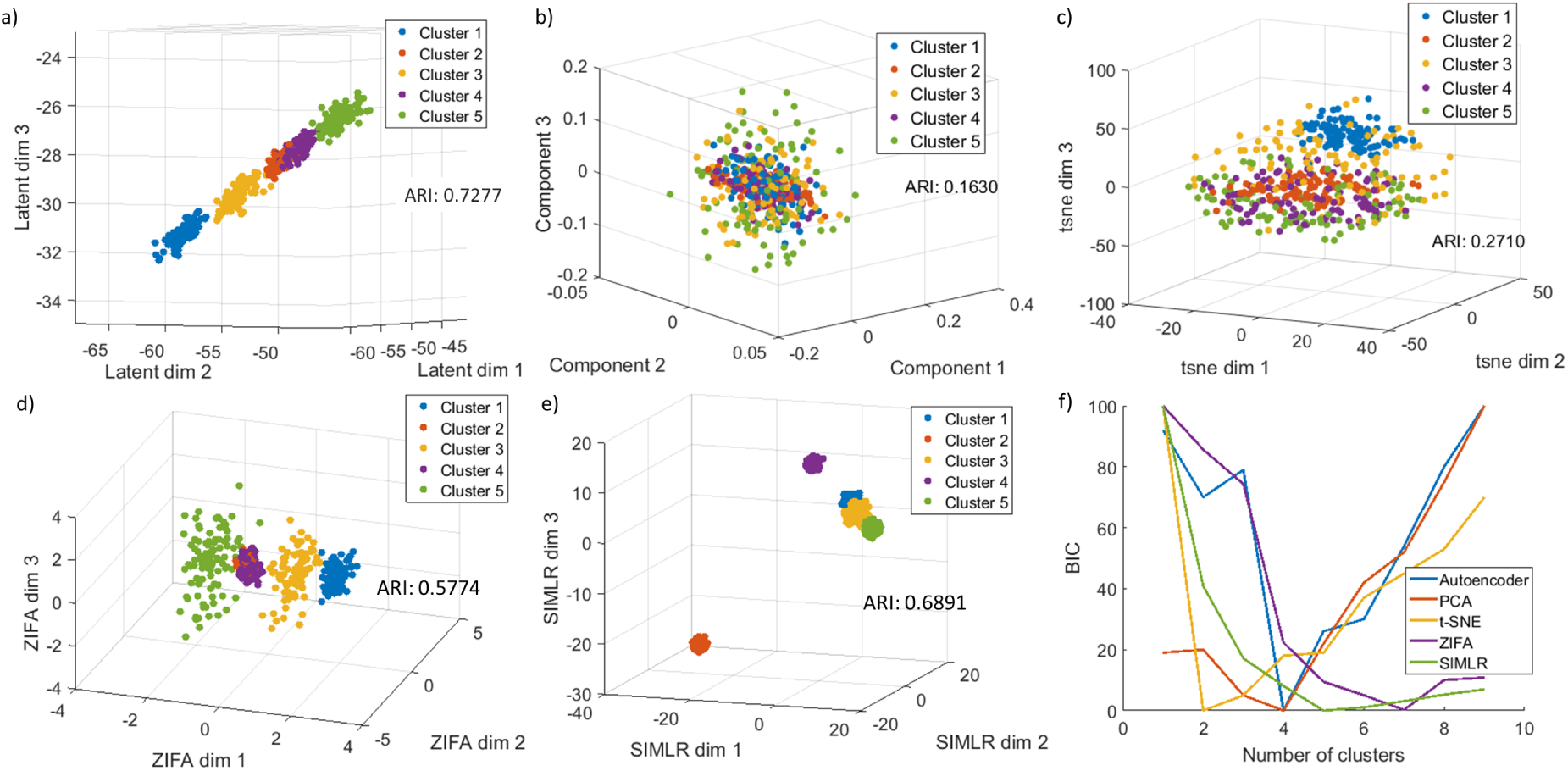
Comparison of the Dhaka method with t-SNE, PCA, ZIFA, and SIMLR on simulated dataset with 2500 completely noisy genes (83% of total genes) without any cluster specific expression. a) Autoencoder, b) PCA, c) t-SNE, d) ZIFA, e) SIMLR. The colors correspond to the ground truth cluster ids. f) Plot of BIC calculated from fitting Gaussian Mixture Model to the 3D projection of the data to estimate number of clusters. The number with lowest BIC is considered as the estimated number of clusters in the data.

We have also compared three different structures of the autoencoder (structure 1: *Input* → 1024 *nodes* → 512 *nodes* → 256 *nodes* → 3 *latent dims*, structure 2: *Input* → 1024 *nodes* → 3 *latent dims*, and 3 *latent dims* in terms of ARI and runtime (Table 1) on the simulated data. The VAE structure 1 (Fig. 1) gives the best ARI score. When we reduce the number of intermediate layers to 1, we see that the runtime decreases slightly but the ARI also decreases from 0.73 to 0.5. We have also tested the effect of increasing the number of intermediate layers to 5.

**Table 1.**
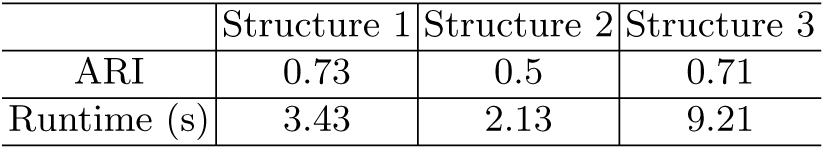
Comparison between structures of autoencoders. Python 3.5, 32GB RAM, 3.4GHz Windows

We see that increasing the number of layers increases the runtime significantly without improving the ARI score. Hence, we used the proposed structure 1 in all of our analysis. We have also compared the runtime with other competing methods, PCA, t-SNE, ZIFA, and SIMLR (See Appendix 1.2, Table S2). We see that only PCA is faster than the proposed method but has very poor ARI score (0.16) compared to Dhaka (0.73).

### Gene expression data

We have next tested the method on four single cell RNA-seq tumor datasets: i) Oligodendroglioma Tirosh et al. (2016b), ii) Glioblastoma Patel et al. (2014), iii) Melanoma Tirosh et al. (2016a), and iv) Astrocytoma Venteicher et al. (2017). We discuss the first three below and the fourth in the appendix.

### Analysis of Oligodendroglioma data

Oligodendrogliomas are a type of glioma that originate from the oligodendrocytes of the brain or from a glial precursor cell. In the Oligodendroglioma dataset the authors profiled six untreated Oligodendroglioma tumors resulting in 4347 cells and 23K genes. The dataset is comprised of both malignant and non-malignant cells. Copy number variations (CNV) were estimated from the *log*2 transformed transcript per million (TPM) RNA-seq expression data. The authors then computed two metrics, lineage score and differentiation score by comparing pre-selected 265 signature genes’ CNV profile for each cell with that of a control gene set. Based on these metrics, the authors determined that the malignant cells are composed from two subpopulations, oligo-like and astro-like, and that both share a common lineage. The analysis also determined the differentiation state of each cell.

Here and in all the following RNA-seq datasets we are using the *log*2 TPM RNA-seq expression data directly skipping the CNV analysis. With only three latent dimensions our algorithm successfully separated malignant cells from non-malignant microglia/macrophage cells (Fig. 3a). We next analyzed the malignant cells only using their relative expression profile (see appendix 1.3), to identify the different subpopulations and the relationship between them. Fig. 3b-c show the projected autoencoder output, where we see two distinct subpopulations originating from a common lineage, thus recapitulating the finding of the original paper. The autoencoder was not only able to separate the two subpopulations, but also to uncover their shared glial lineage. To compare the results with the original paper, we have plotted the scatter plot with color corresponding to lineage score (Fig. 3b) and differentiation score (Fig. 3c) from Tirosh et al. (2016b). We can see from the figure that the autoencoder can separate oligo-like and astro-like cells very well by placing them in opposite arms of the v-structure. In addition, figure Fig 3c shows that most of the cells with stem like property are placed near the bifurcation point of the v-structure. From the figure it looks like Latent dim 1 and 2 correlates with the lineage score whereas Latent dim 3 correlates with differentiation. However, since VAEs are stochastic in nature, there is no guarantee that same latent dimension will always correlate with the same score unlike PCA. Although Dhaka can consistently capture the v-structures, the correspondence between the latent dimensions and lineage/differentiation score might change from one run to the next (see Supporting Fig. S2).

**Fig. 3.**
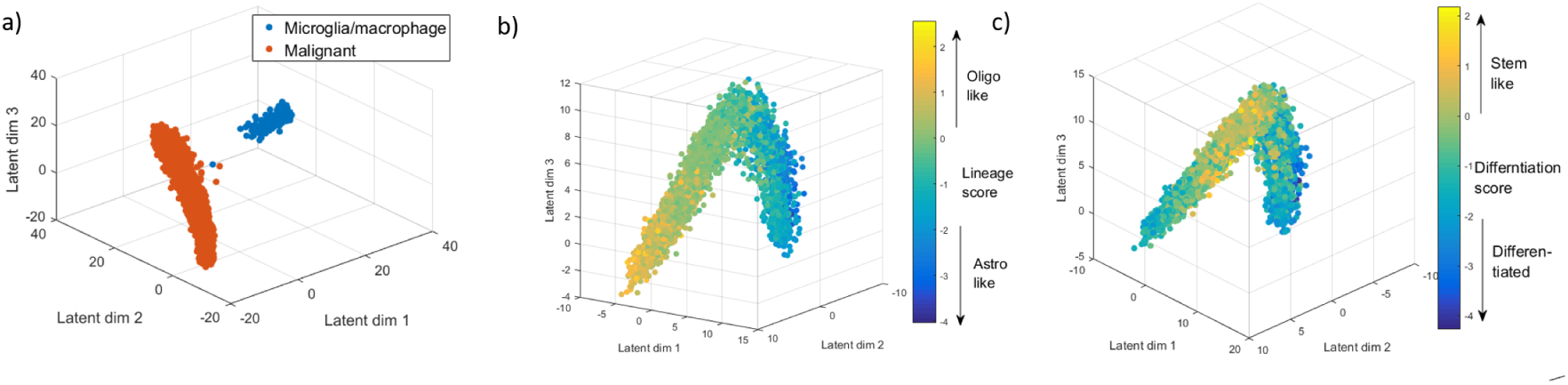
Oligodendroglioma dataset.a) Autoenocder projection separating malignant cells from non-malignant microglia/macrophage cells. b)-c) Autoencoder output from relative expression profile of malignant cells using 265 signatures genes. b) Each cell is colored by their assigned lineage score which differentiates the oligo-like and astro-like subpopulations. c) Each cell is colored by their assigned differentiation score, which shows that most stem like cells are indeed placed near the bifurcation point.

The analysis discussed above was based on the 265 signature genes that were reported in the original paper. We next tested whether a similar structure can be learned from auto selected genes, instead of using these signature genes. Malignant and non-malignant cells were clearly separated in this scenario too (Supporting Fig. S3). Fig. 4a shows the autoencoder projection of the malignant cells only using 5000 auto-selected genes based on *Ā* score (see appendix 1.1). As we can see from Fig. 4a, the autoencoder can learn similar structure without the need for supervised prior knowledge. We also compared the autoencoder output for this data to PCA, t-SNE, ZIFA, and SIMLR (Fig. 4b-e). As can be seen, PCA and ZIFA can separate the oligo-like and astro-like structure to some extent, but their separation is not as distinct as the autoencoder output. On the other hand, t-SNE and SIMLR can recover clusters of cells from the same tumor but completely fails to identify the underlying lineage and differentiation structure of the data. To quantify how well the lineage and differentiation metrics are preserved in the projections we have computed Spearman rank correlation score Zar (1998) of the scoring metrics (Lineage and differentiation scores) with the projections of the autoencoder and other comparing methods. Since the ground truth is a 2D metric, we computed correlation with 2D projections from Dhaka and other competing methods (see supporting Fig. S4a for 2D projection from Dhaka). From the correlation scores we can clearly see that Dhaka performs significantly better than the other methods. We have also computed and compared the correlation score on the 265 signature gene scenario (see Supporting fig. S5). With using only signature genes the correlation score from Dhaka is 0.76, whereas the nearest competing methods t-SNE and SIMLR scores 0.57 and 0.52, respectively. Method of correlation score computation can be found in Appendix 1.4.

**Fig. 4.**
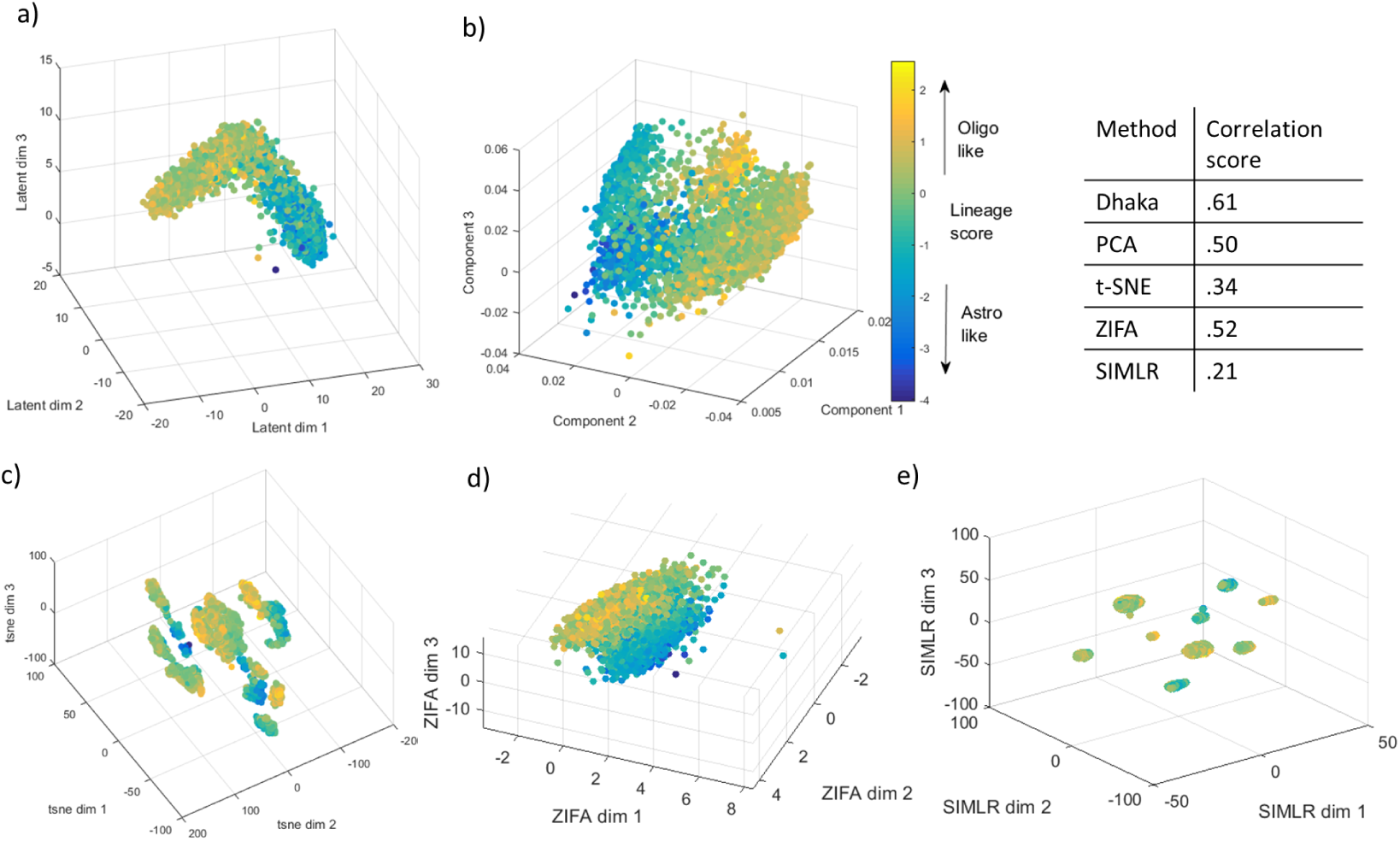
Comparison of variational autoencoder with PCA, t-SNE, ZIFA, and SIMLR on Oligodendroglioma dataset with 5000 autoselected genes. a) Autoencoder, b) PCA, c) t-SNE, d) ZIFA, e) SIMLR projections colored by the lineage score. The Spearman rank correlation scores of the scoring metric (lineage and differentiation score) and the learned projections are shown in tabular form. We can clearly see that Dhaka preserves the original scoring metric the best.

#### Robustness analysis

A key issue with the analysis of scRNA-Seq data is dropout. In scRNA-Seq data we often see transcripts that are not detected even though the particular gene is expressed, which is known as the ‘dropout’. This happens mostly because of the low genomic quantity used for scRNA-Seq. We have tested the robustness of the autoencoder to dropouts in the Oligodendroglioma dataset. We tested several different dropout percentages ranging from 0 to 50% (Supporting Fig. S6a). Supporting Fig. S6c, e,g shows the histogram of dropout fractions of the genes in the dataset after artificially forcing 20%, 30%, and 50% more genes to be dropped out. Note that we cannot go beyond 50% in our analysis since several genes are already zero in the original data. Supporting Fig. S6b,d,f,h shows the projection of the autoencoder after adding 0%, 20%, 30%, and 50% more dropout genes, respectively. We observe that when the additional dropout rate is 30% or less, the autoencoder can still retain the v-structure even though the cells are a bit more dispersed. At 50% we lose the v-structure, but the method can still separate oligo-like and astro-like cells even with this highly sparse data.

#### Analysis of marker genes in the Oligodendroglioma dataset

We further investigated the autoencoder learned structure to discover genes that are correlated with the lineages. To obtain trajectories for genes in the two lineages of the Oligodendroglioma dataset, we first segmented the autoencoder projected output into 9 clusters using Gaussian mixture model (Fig 5). Clusters 1 to 4 correspond to the oligo branch and clusters −4 to −1 correspond to the astro branch, while cluster 0 represents the bifurcation point. The choice to divide the cells into 9 clusters is arbitrary to show the difference between the two branches. For each gene we compute two profiles, one for the astro branch and one for the oligo branch. We select the genes that have statistically significant differential expression profile in astro branch and oligo branch using Bonferroni corrected t-test. With bonferroni corrected p-value < 0.05, we find 1197 DE genes among 23K original genes. We have also separately identified genes that are up regulated and down regulated in the two lineages (see list of genes in the supporting website). Expression profiles of a few of these genes are shown in Fig. 5b)-e). Many of the genes we identified were known to be related to other types of cancers or neurological disorders, but so far have not been associated with Oligodendroglioma. For example, TFG which is upregulated in the oligo-branch was previously affiliated in neuropathy Ishiura et al. (2012). *DDX39B* gene is not directly related to cancer but is found to be localized near genes encoding for tumor necrosis factor *α* and β Kikuta et al. (2012). Both *HEXB* and *RGMA* genes are up regulated in the astro-branch. These genes were previously identified in neurological disorders such as Sandhoff disease Redonnet-Vernhet et al. (1996) and multiple sclerosis Nohra et al. (2010), respectively. Our analysis suggests that they are key players in the Oligodendroglioma pathway as well.

**Fig. 5.**
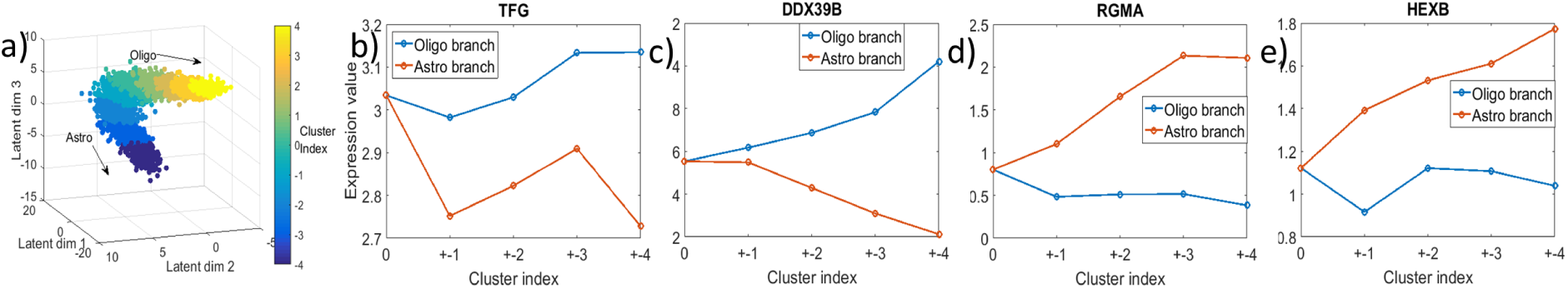
New gene markers for astro-like and oligo-like lineages. a) Segmenting autoencoder projected output to 9 clusters. Clusters −4, −3, −2, −1 belongs to astro branch and clusters 1,2,3,4 belong to oligo branch. Cluster 0 represent the origin of bifurcation. b)-e) Expression profiles of couple of the top differntially expressed genes in the two lineages. b)-c) upregulated in the oligo-branch, d)-e) upregulated in the astro-branch.

### Analysis of Glioblastoma data

The next dataset we looked at is the Glioblastoma dataset Patel et al. (2014). This dataset contains 420 malignant cells with ~ 6000 expressed genes from six tumors. In this relatively small cohort of cells the authors did not find multiple subpopulations. However, they identified a stemness gradient across the cells from all six tumors Patel et al. (2014), meaning the cells gradually evolve from a stemlike state to a more differentiated state. When we applied the Dhaka autoencoder to the expression profiles of the malignant cells, the cells were arranged in a chain like structure Fig. 6a). To correlate the result with the underlying biology, we computed stemness score from the signature genes reported in the original paper (78 genes in total) Patel et al. (2014). The score is computed as the ratio of average expression of the stemness signature genes to the average expression of all remaining genes Patel et al. (2014). When we colored the scattered plot according to the corresponding stemness score of each cell, we see a chain like evolutionary structure where cells are gradually progressing form a stemlike state to a more differentiated state. As before, PCA, t-SNE, ZIFA, and SIMLR projections (Fig. 6b-e), fail to capture the underlying structure of this differentiation process. We do see some outliers, blue dots around the yellow dots in Fig. 6a. These outliers are also evident in ZIFA and SIMLR projections (blue dots). However, when we quantify the correlation of the autoencoder projection with the stemness score we can see that it clearly outperforms the other competing methods despite the outliers. After learning the structures we also wanted to see whether we can identify new marker genes for the stemness to differentiated program. For this, we reduced the latent dimension to 1 (since we see almost linear projection). Next, we computed Spearman rank correlation Zar (1998) of the 1D projection with every gene in the dataset. We have plotted a few of the top ranked positive (up regulated in the stemlike cells) and negative correlated genes (down regulated in the stemlike cells) (Fig. 7a). Despite the noisy expression profile, we do see a clear trend when a line is fitted (red). Among the discovered markers, *TPT1* was identified as one of the key tumor proteins Arcuri et al. (2004). Both *RPS27* and *TPT1* were found to be significant in other forms of cancer, such as Melanoma Dai et al. (2010) and prostate cancer Arcuri et al. (2004) and our results indicate that they may be involved in Glioblastoma as well. Among the downregulated genes, *CLU* was identified in the original paper Patel et al. (2014) to be affiliated in Glioblastoma pathway whereas *CANX* was previously not identified as a marker for Glioblastoma. A complete list of correlated marker genes can be found in the supporting website.

**Fig. 6.**
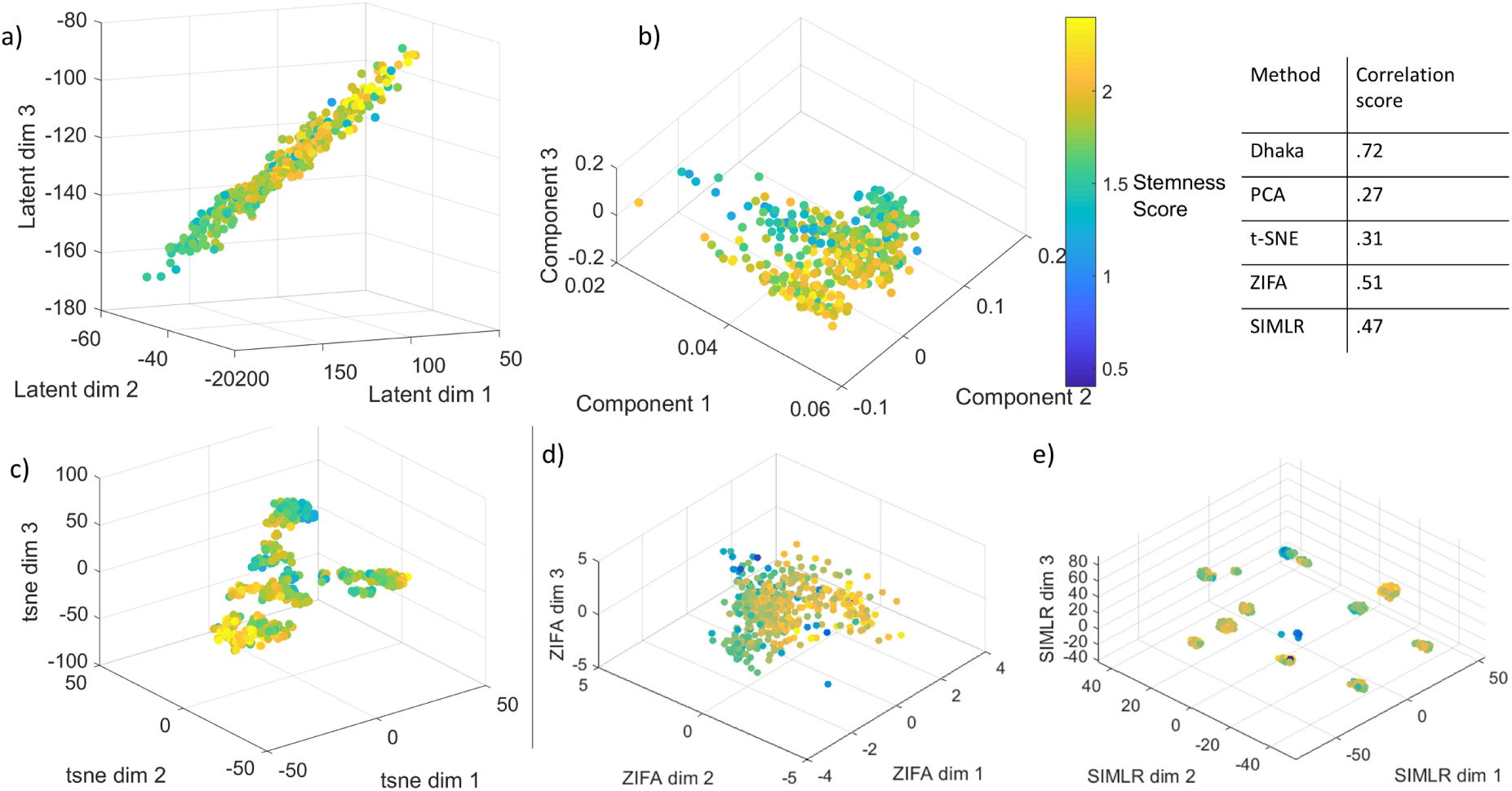
Comparison of variational autoencoder with PCA, t-SNE, ZIFA, and SIMLR on Glioblastoma dataset. a) Autoencoder, b) PCA, c) t-SNE, d) ZIFA, e) SIMLR projections colored by stemness score. The Spearman rank correlation scores of the stemness score and the learned projections are shown in tabular form. We can clearly see that Dhaka preserves the original stemness score the best.

**Fig. 7.**
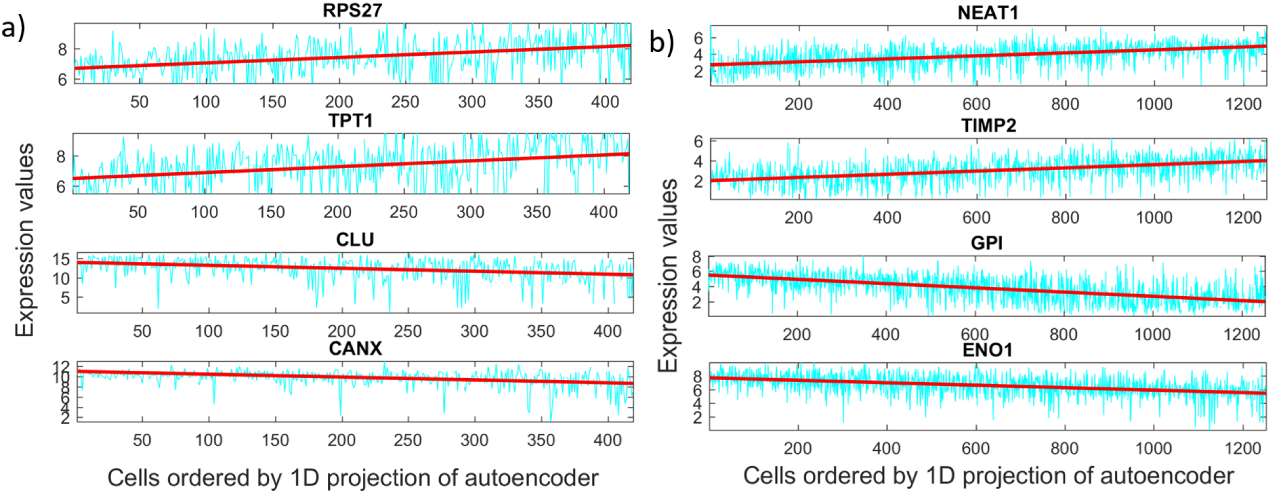
New marker gene: a) Glioblastoma stemness program b) Melanoma MITF-AXL program.

### Analysis of Melanoma data

The Melanoma cancer dataset Tirosh et al. (2016a) profiled 1252 malignant cells with ~ 23K genes from 19 samples. The expression values are log2 transformed transcript per million. When we used the relative expression values of 5000 auto-selected genes (based on *Ā* score) to the Dhaka autoencoder we saw two very distinct clusters of cells, revealing the intra-tumor heterogeneity of the Melanoma samples (Fig. 8a). In the original paper, the authors identified two expression programs related to *MITF* and *AXL* genes that give rise to a subset of cells that are less likely to respond to targeted therapy. The signature score for these programs were calculated by identifying genesets correlated with these two programs. The authors identified a total of 200 signature genes. We computed *MITF-AXL* signature score by computing the ratio of average expression of the signature genes and average expression of all remaining genes. When we colored the scattered plot with the *MITF-AXL* score, we indeed see that the clusters correspond to the *MITF-AXL* program, with one cluster scoring high and the other scoring low for these signature genes. Again, as can be seen from the figures and the correlation scores, such heterogeneity is not properly captured by t-SNE, PCA, ZIFA, and SIMLR (Fig 8b-e).

**Fig. 8.**
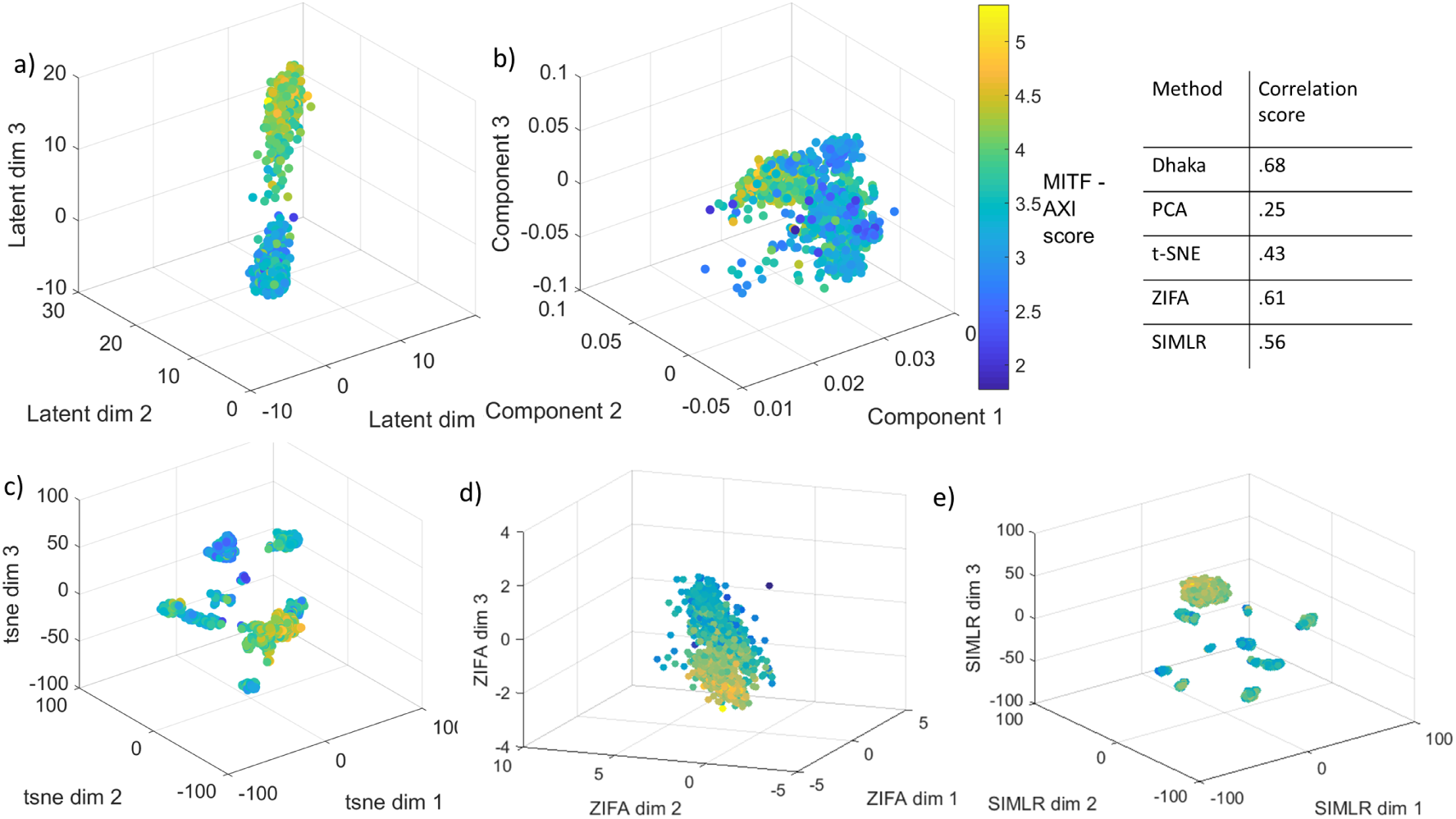
Comparison of variational autoencoder with PCA, t-SNE, ZIFA, and SIMLR on Melanoma dataset. a) Autoencoder, b) PCA, c) t-SNE, d) ZIFA, e) SIMLR projections colored by MITF-AXL score. The Spearman rank correlation scores with the scoring metric (MITF-AXL score) and the autoencoder and the reported method projections are shown in tabular form. We can clearly see that Dhaka preserves the original MITF-AXL score the best.

For this case too, we see almost a linear projection. To find new gene markers, we again computed 1D latent projection of the single cells and computed gene correlation. We have plotted a set of new marker genes both up and down regulated (Fig. 7b)). The *NEAT1* is a non-coding RNA, which acts as a transcriptional regulator for numerous genes, including some genes involved in cancer progression Geirsson et al. (2003). *TIMP2* gene plays a critical role in suppressing proliferation of endothelial cells and now we can see it is also relevant in the Melanoma cells Vairaktaris et al. (2009). Among the down regulated genes, *GPI* functions as tumor-secreted cytokine and an angiogenic factor, which is very relevant to any cancer progression Funasaka et al. (2001). The last correlated down regulated gene *ENO1* is also known as tumor suppressor Abu-Odeh et al. (2014). We have also looked whether the projection can recover some known gene marker dynamics or not. Four of the known gene markers are plotted in supporting Fig. S7 (in Appendix). A complete set of gene markers can be found in the supporting website.

We also investigated whether Dhaka can capture the same trends in 2D projections as well. We have computed 2D projections of Oligodendroglioma, Glioblastoma, and Melanoma (Supporting Fig. S4). The structures we see in the 3D projections are retained in the 2D projections as well. However, the correlation score decreases slightly for Glioblastoma and Melanoma indicating that a 3D projection may be more informative in capturing the underlying biology.

Monocle Trapnell et al. (2014) is a another popular single-cell RNA-seq tool. Even though the scope for Monocle and Dhaka is different (Monocle finds pseudotime ordering where Dhaka is a dimensionality reduction method), we have further compared Dhaka to pseudotime ordering for the RNA-Seq data. As expected, Monocle was unable to recover the structure of the tumor subpopulation (with correlation score of just 0.32 (Oligodendroglioma), 0.23 (Glioblastoma), and 0.27 (Melanoma)). See Supporting Fig. S9 for complete results.

We have also analyzed another scRNA-seq tumor dataset, Astrocytoma. Due to space constraint we have moved the analysis to the Appendix.

### Copy number variation data

To test the generality of the method we also tested Dhaka with copy number variation data. We used copy number profiles from two xenograft breast tumor samples (xenograft 3 and 4, representing two consecutive time points) Zahn et al. (2017). 260 cells were profiled from xenograft 3 and 254 from xenograft 4. Both of these datasets have around 20K genomic bin count. Cells were sequenced at a very low depth of 0.05X which results in noisy profiles. Copy numbers were estimated using a hidden Markov model Wang et al. (2007). When we analyzed the copy number profile for xenograft 3, the autoencoder identified 1 major cluster of cells and 1 minor cluster of cells (Fig 9a). The identified clusters agree with the phylogenetic reconstruction analysis in the original paper. Fig. 9b shows the copy number profiles of cells organized by phylogenetic analysis. Even though the copy number profiles are mostly similar in most parts of the genome, we do see that there is a small number of cells that have two copies (as opposed to one in the majority of cells, marked by red circle) in the x chromosome. The autoencoder was able to correctly differentiate the minor cluster of cells from the rest. Next we analyzed the xenograft 4 samples. The projected autoencoder output showed only 1 cluster which overlaps the major cluster identified for xenograft 3. We believe that the minor cluster from xenograft 3 probably did not progress further after serial passaging to the next mouse, whereas the major cluster persisted. This observation also agrees with the claim stated in the original paper Zahn et al. (2017) that after serial passaging only one cluster remained in xenograft 4 which is a descendant of the major cluster in xenograft 3.

**Fig. 9.**
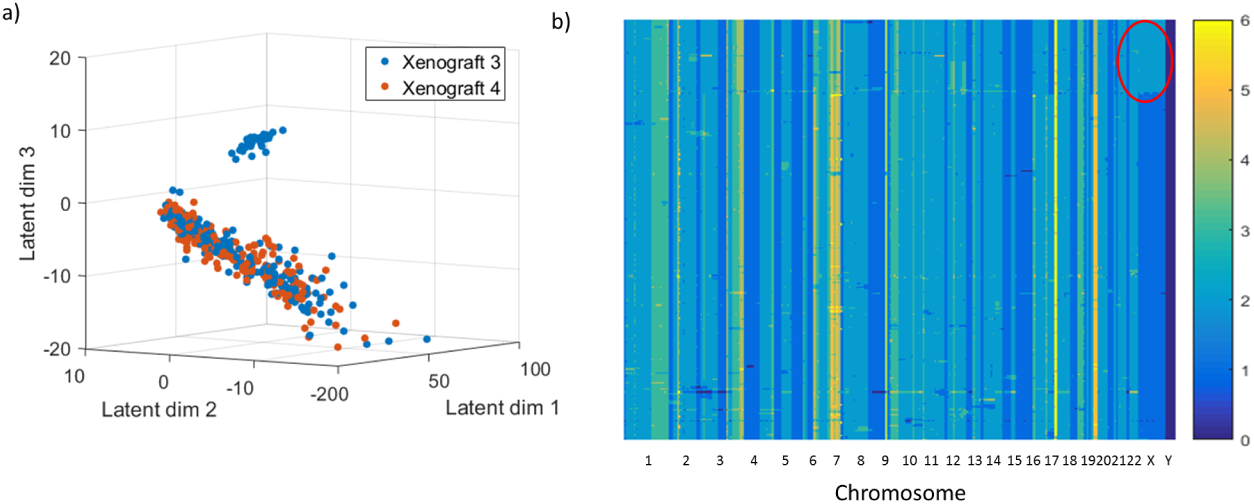
Autoencoder ouptut of two xenograft breast tumor samples’ copy number profile. a) Identification of two subpopulations of cells in xenograft 3 and one subpopulation in xenograft 4. b) Copy number profile of cells in xenograft 3 ordered by phylogenetic analysis, which shows that there are indeed two groups of cells present in the data.

## Discussion

In this paper, we have proposed a new way of extracting useful features from single cell genomic data. We have shown that Dhaka improves upon prior methods for the analysis of scRNA-Seq data, and that, unlike some of these prior methods Pierson and Yau (2015), Wang et al. (2017), it can be applied to other types of data including copy number variations. The method is completely unsupervised and requires minimal preprocessing of the data. Using our method we were able to reconstruct lineage and differentiation ordering for several single cell tumor samples. The autoencoder successfully separated oligo-like and astro-like cells along with their differentiation status for Oligodendroglioma scRNA-Seq data and has also successfully captured the differentiation trajectory of Glioblastoma cells. Similar results were obtained for Melanoma and Astrocytoma. The autoencoder projections have also revealed several new marker genes for the cancer types analyzed. When applied to copy number variation data the method was able to identify heterogeneous tumor populations for breast cancer samples. in future, we will investigate larger single cell copy number datasets with more cells and more subpopulations.

An advantage of the autoencoder method is its ability to handle dropouts. Several single cell algorithms require preprocessing to explicitly model the dropout rates. As we have shown, our method is robust and can handle very different rates eliminating the need to estimate this value.

While Dhaka leads to better ordering of the different cell lineages, it may be harder to directly interpret the meaning of the different dimensions in the Dhaka based embedding. Unlike the deterministic PCA, VAEs are stochastic in nature and so there is no guarantee that same latent dimension will always correlate with the same score or that the first dimension would always represent the same biological function. While this is indeed a drawback of Dhaka, for the data analyzed in this paper we consistently see that Dhaka results are more in line with known biology when compared to the deterministic methods.

While our focus here was primarily on the identification of sub populations and visualization, the latent representation generated by the autoencoder could be used in pseudotime ordering algorithms as well Trapnell et al. (2014), Setty et al. (2016). These methods often rely on t-SNE/PCA as the first step and replacing these with the autoencoder is likely to yield more accurate results. The variational autoencoder does not only cluster the cells, it can also represent an evolutionary trajectory, for example, the V structure for the Oligodendroglioma. Hence it can also be useful in phylogenetic analysis. Potential future work would focus on using this to identify key genes that align with the progression and mutations that help drive the different populations.

## Bibliography

Mohammad Abu-Odeh, Tomer Bar-Mag, Haiming Huang, TaeHyung Kim, Zaidoun Salah, Suhaib K Abdeen, Marius Sudol, Dana Reichmann, Sachdev Sidhu, Philip M Kim, et al. Characterizing ww domain interactions of tumor suppressor wwox reveals its association with multiprotein networks. Journal of Biological Chemistry, 289(13):8865–8880, 2014.

Noemi Andor, Trevor A Graham, Marnix Jansen, Li C Xia, C Athena Aktipis, Claudia Petritsch, Hanlee P Ji, and Carlo C Maley. Pan-cancer analysis of the extent and consequences of intratumor heterogeneity. Nature medicine, 22(1):105, 2016.

Felice Arcuri, Stefania Papa, Antonietta Carducci, Roberta Romagnoli, Sabrina Liberatori, Maria Giovanna Riparbelli, Jean-Charles Sanchez, Piero Tosi, and Maria Teresa del Vecchio. Translationally controlled tumor protein (tctp) in the human prostate and prostate cancer cells: expression, distribution, and calcium binding activity. The Prostate, 60(2):130–140, 2004.

Yuemeng Dai, Spencer E Pierson, W Dudney, and Brendan C Stack. Extraribosomal function of metallopanstimulin-1: reducing paxillin in head and neck squamous cell carcinoma and inhibiting tumor growth. International journal of cancer, 126(3):611–619, 2010.

Elza C de Bruin, Nicholas McGranahan, Richard Mitter, Max Salm, David C Wedge, Lucy Yates, Mariam Jamal-Hanjani, Seema Shafi, Nirupa Murugaesu, Andrew J Rowan, et al. Spatial and temporal diversity in genomic instability processes defines lung cancer evolution. Science, 346 (6206):251–256, 2014.

David DeTomaso and Nir Yosef. Fastproject: a tool for low-dimensional analysis of single-cell rna-seq data. BMC bioinformatics, 17(1):315, 2016.

Jean Fan, Neeraj Salathia, Rui Liu, Gwendolyn E Kaeser, Yun C Yung, Joseph L Herman, Fiona Kaper, Jian-Bing Fan, Kun Zhang, Jerold Chun, et al. Characterizing transcriptional heterogeneity through pathway and gene set overdispersion analysis. Nature methods, 13(3):241, 2016.

Tatsuyoshi Funasaka, Arayo Haga, Avraham Raz, and Hisamitsu Nagase. Tumor autocrine motility factor is an angiogenic factor that stimulates endothelial cell motility. Biochemical and biophysical research communications, 284(5):1116–1125, 2001.

Charles Gawad, Winston Koh, and Stephen R Quake. Single-cell genome sequencing: current state of the science. Nature reviews. Genetics, 17(3):175, 2016.

Arnar Geirsson, Raymond J Lynch, Indu Paliwal, Alfred LM Bothwell, and Graeme L Hammond. Human trophoblast noncoding rna suppresses ciita promoter iii activity in murine b-lymphocytes. Biochemical and biophysical research communications, 301(3):718–724, 2003.

Alice Giustacchini, Supat Thongjuea, Nikolaos Barkas, Petter S Woll, Benjamin J Povinelli, Christopher AG Booth, Paul Sopp, Ruggiero Norfo, Alba Rodriguez-Meira, Neil Ashley, et al. Single-cell transcriptomics uncovers distinct molecular signatures of stem cells in chronic myeloid leukemia. Nature medicine, 23(6):692–702, 2017.

Aman Gupta, Haohan Wang, and Madhavi Ganapathiraju. Learning structure in gene expression data using deep architectures, with an application to gene clustering. In Bioinformatics and Biomedicine (BIBM), 2015 IEEE International Conference on, pages 1328–1335. IEEE, 2015.

Hiroyuki Ishiura, Wataru Sako, Mari Yoshida, Toshitaka Kawarai, Osamu Tanabe, Jun Goto, Yuji Takahashi, Hidetoshi Date, Jun Mitsui, Budrul Ahsan, et al. The trk-fused gene is mutated in hereditary motor and sensory neuropathy with proximal dominant involvement. The American Journal of Human Genetics, 91(2):320–329, 2012.

Ian T Jolliffe. Principal component analysis and factor analysis. In Principal component analysis, pages 115–128. Springer, 1986.

James M Joyce. Kullback-leibler divergence. In International Encyclopedia of Statistical Science, pages 720–722. Springer, 2011.

Kazutaka Kikuta, Daisuke Kubota, Tsuyoshi Saito, Hajime Orita, Akihiko Yoshida, Hitoshi Tsuda, Yoshiyuki Suehara, Hitoshi Katai, Yasuhiro Shimada, Yoshiaki Toyama, et al. Clinical proteomics identified atp-dependent rna helicase ddx39 as a novel biomarker to predict poor prognosis of patients with gastrointestinal stromal tumor. Journal of proteomics, 75(4):1089–1098, 2012.

Diederik P Kingma and Max Welling. Auto-encoding variational bayes. *arXiv preprint arXiv*:1312.6114, 2013.

Yann LeCun, Yoshua Bengio, and Geoffrey Hinton. Deep learning. Nature, 521(7553):436–444, 2015.

Xiangyu Li, Weizheng Chen, Yang Chen, Xuegong Zhang, Jin Gu, and Michael Q Zhang. Network embedding-based representation learning for single cell rna-seq data. Nucleic acids research, 45(19):e166-e166, 2017.

Chieh Lin, Siddhartha Jain, Hannah Kim, and Ziv Bar-Joseph. Using neural networks for reducing the dimensions of single-cell rna-seq data. Nucleic Acids Research, 2017.

Laurens van der Maaten and Geoffrey Hinton. Visualizing data using t-sne. Journal of Machine Learning Research, 9(Nov):2579–2605, 2008.

Jae-Woong Min, Woo Jin Kim, Jeong A Han, Yu-Jin Jung, Kyu-Tae Kim, Woong-Yang Park, Hae-Ock Lee, and Sun Shim Choi. Identification of distinct tumor subpopulations in lung adenocarcinoma via single-cell rna-seq. PloS one, 10(8):e0135817, 2015.

Nicholas E Navin and James Hicks. Tracing the tumor lineage. Molecular oncology, 4(3):267–283, 2010.

R Nohra, AD Beyeen, JP Guo, M Khademi, E Sundqvist, MT Hedreul, F Sellebjerg, C Smestad, AB Oturai, HF Harbo, et al. Rgma and il21r show association with experimental inflammation and multiple sclerosis. Genes and immunity, 11(4):279–293, 2010.

Anoop P Patel, Itay Tirosh, John J Trombetta, Alex K Shalek, Shawn M Gillespie, Hiroaki Wakimoto, Daniel P Cahill, Brian V Nahed, William T Curry, Robert L Martuza, et al. Single-cell rna-seq highlights intratumoral heterogeneity in primary glioblastoma. Science, 344(6190):1396–1401, 2014.

Emma Pierson and Christopher Yau. Zifa: Dimensionality reduction for zero-inflated single-cell gene expression analysis. Genome biology, 16(1):241, 2015.

Isabelle Redonnet-Vernhet, Don J Mahuran, Robert Salvayre, Frederic Dubas, and Thierry Levade. Significance of two point mutations present in each hexb allele of patients with adult gm2 gangliosidosis (sandhoff disease) homozygosity for the ile207 val substitution is not associated with a clinical or biochemical phenotype. Biochimica et Biophysica Acta (BBA)-Molecular Basis of Disease, 1317(2):127–133, 1996.

Sam T Roweis and Lawrence K Saul. Nonlinear dimensionality reduction by locally linear embedding. science, 290(5500):2323–2326, 2000.

Hege G Russnes, Nicholas Navin, James Hicks, and Anne-Lise Borresen-Dale. Insight into the heterogeneity of breast cancer through next-generation sequencing. The Journal of clinical investigation, 121(10):3810, 2011.

Manu Setty, Michelle D Tadmor, Shlomit Reich-Zeliger, Omer Angel, Tomer Meir Salame, Pooja Kathail, Kristy Choi, Sean Bendall, Nir Friedman, and Dana Pe’er. Wishbone identifies bifurcating developmental trajectories from single-cell data. Nature biotechnology, 34(6):637–645, 2016.

Tijmen Tieleman and Geoffrey Hinton. Lecture 6.5-rmsprop: Divide the gradient by a running average of its recent magnitude. COURSERA: Neural networks for machine learning, 4(2):26–31, 2012.

Itay Tirosh, Benjamin Izar, Sanjay M Prakadan, Marc H Wadsworth, Daniel Treacy, John J Trombetta, Asaf Rotem, Christopher Rodman, Christine Lian, George Murphy, et al. Dissecting the multicellular ecosystem of metastatic melanoma by single-cell rna-seq. Science, 352(6282): 189–196, 2016a.

Itay Tirosh, Andrew S Venteicher, Christine Hebert, Leah E Escalante, Anoop P Patel, Keren Yizhak, Jonathan M Fisher, Christopher Rodman, Christopher Mount, Mariella G Filbin, et al. Single-cell rna-seq supports a developmental hierarchy in human oligodendroglioma. Nature, 539(7628):309–313, 2016b.

Cole Trapnell, Davide Cacchiarelli, Jonna Grimsby, Prapti Pokharel, Shuqiang Li, Michael Morse, Niall J Lennon, Kenneth J Livak, Tarjei S Mikkelsen, and John L Rinn. The dynamics and regulators of cell fate decisions are revealed by pseudotemporal ordering of single cells. Nature biotechnology, 32(4):381–386, 2014.

E Vairaktaris, Z Serefoglou, D Avgoustidis, C Yapijakis, E Critselis, A Vylliotis, S Spyridonidou, S Derka, S Vassiliou, E Nkenke, et al. Gene polymorphisms related to angiogenesis, inflammation and thrombosis that influence risk for oral cancer. Oral oncology, 45(3):247–253, 2009.

Andrew S Venteicher, Itay Tirosh, Christine Hebert, Keren Yizhak, Cyril Neftel, Mariella G Filbin, Volker Hovestadt, Leah E Escalante, McKenzie L Shaw, Christopher Rodman, et al. Decoupling genetics, lineages, and microenvironment in idh-mutant gliomas by single-cell rna-seq. Science, 355(6332):eaai8478, 2017.

Bo Wang, Junjie Zhu, Emma Pierson, Daniele Ramazzotti, and Serafim Batzoglou. Visualization and analysis of single-cell rna-seq data by kernel-based similarity learning. Nature Methods, 14(4):414–416, 2017.

Kai Wang, Mingyao Li, Dexter Hadley, Rui Liu, Joseph Glessner, Struan FA Grant, Hakon Hakonarson, and Maja Bucan. Penncnv: an integrated hidden markov model designed for high-resolution copy number variation detection in whole-genome snp genotyping data. Genome research, 17 (11):1665–1674, 2007.

Chen Xu and Zhengchang Su. Identification of cell types from single-cell transcriptomes using a novel clustering method. Bioinformatics, 31(12):1974–1980, 2015.

Xianxue Yu, Guoxian Yu, and Jun Wang. Clustering cancer gene expression data by projective clustering ensemble. PloS one, 12(2):e0171429, 2017.

Hans Zahn, Adi Steif, Emma Laks, Peter Eirew, Michael VanInsberghe, Sohrab P Shah, Samuel Aparicio, and Carl L Hansen. Scalable whole-genome single-cell library preparation without preamplification. Nature methods, 14(2):167–173, 2017.

Jerrold H Zar. Spearman rank correlation. Encyclopedia of Biostatistics, 1998.

Chenghang Zong, Sijia Lu, Alec R Chapman, and X Sunney Xie. Genome-wide detection of singlenucleotide and copy-number variations of a single human cell. Science, 338(6114):1622–1626, 2012.

